# Stenoparib, an inhibitor of cellular poly (ADP-ribose) polymerases (PARPs), blocks *in vitro* replication of SARS-CoV-2 variants

**DOI:** 10.1101/2021.11.03.467186

**Authors:** Katherine E Zarn, Sierra A Jaramillo, Anthony R Zapata, Nathan E Stone, Ashley N Jones, Haley E Nunnally, Erik W Settles, Ken Ng, Paul S Keim, Steen Knudsen, Patricia M Nuijten, Aloys SL Tijsma, Christopher T French

## Abstract

We recently published a preliminary assessment of the activity of a poly (ADP-ribose) polymerase (PARP) inhibitor, stenoparib, also known as 2X-121, which inhibits viral replication by affecting pathways of the host. Here we show that stenoparib effectively inhibits a SARS-CoV-2 *wt* (BavPat1/2020) strain and four additional variant strains; *alpha* (B.1.1.7), *beta* (B.1.351), *delta* (B.1.617.2) and *gamma* (P.1) *in vitro,* with 50% effective concentration (EC_50_) estimates of 4.1 μM, 8.5 μM, 24.1 μM, 8.2 μM and 13.6 μM, respectively. A separate experiment focusing on a combination of 10 μM stenoparib and 0.5 μM remdesivir, an antiviral drug, resulted in over 80% inhibition of the *alpha* (B.1.1.7) variant, which is substantially greater than the effect achieved with either drug alone, suggesting at least additive effects from combining the different mechanisms of activity of stenoparib and remdesivir.

## Introduction

As of March 2022, the coronavirus disease (COVID-19) pandemic, caused by SARS-CoV-2, has caused over 446 million infections and over 6 million deaths worldwide [1]. Although protective vaccines are available, the pandemic continues, and both old and new SARS-CoV-2 variants exhibit varying degrees of disease severity and resistance to vaccination. Additional effective therapeutics are urgently needed. Here we describe the antiviral activity of a small molecule, stenoparib, an inhibitor of mammalian poly (ADP-ribose) polymerases (PARPs). We show that stenoparib effectively inhibits replication of SARS-CoV-2 wild-type (*wt*) and variant strains *in vitro*.

SARS-CoV-2 has undergone adaptation and mutation since the beginning of the COVID-19 pandemic, resulting in new variants of the virus. Only two antiviral drugs, remdesivir and molnupiravir, or treatment with monoclonal antibodies, have been approved by the United States Food and Drug Administration as COVID-19 therapies under the Emergency Use Authorization [2, 3]. Remdesivir and molnupiravir are nucleoside analogs that affect the activity of the RNA-dependent RNA polymerase (RdRp). After incorporation into viral RNA, remdesivir stalls the RdRp during elongation [4], while molnupiravir results in the incorporation of mutations that can be functionally deleterious [5].

We recently published a study on the activity of a poly (ADP-ribose) polymerase (PARP) inhibitor, stenoparib, also known as 2X-121, which inhibits viral replication by affecting pathways of the host [6] as opposed to targeting viral replication. ADP-ribosylation (ADPR) pathways may have either anti- or pro-viral properties, and their importance in host-virus interactions is becoming increasingly recognized [7]. Unlike remdesivir, which inhibits viral replication downstream of entry into the cell, stenoparib inhibits virus entry and post-entry processes [6]. Stenoparib inhibited the SARS-CoV-2 USA-WA1/2020 virus and the HCoV-NL63 human seasonal respiratory coronavirus *in vitro*, exhibiting dose-dependent suppression of virus multiplication and cell-cell spread in cell culture [6]. Stenoparib exhibits a unique dual activity against the PARP1, 2 and PARP5a, 5b (tankyrase 1, 2) enzymes, which are important intermediates in the Wnt/β-catenin immune checkpoint [8, 9]. As a host-targeting therapeutic that does not directly select for resistance in viruses, we hypothesize that stenoparib should be able to inhibit all SARS-CoV-2 variant strains.

## Materials and Methods

### Cell Culture

For the ViroSpot reduction assays, the antiviral activity of stenoparib on SARS-CoV-2 was assessed *in vitro* using Vero E6 *Cercopithecus aethiops* kidney cells (cat # CRL-1586, ATCC, Manassas, VA, USA) maintained in Dulbecco’s Modified Eagle Medium (DMEM; cat #BE12-733F, Lonza, Basel, Switzerland) supplemented with 2 mM L-glutamine (cat # BE17-605E, Lonza, Basel, Switzerland), 100 U/ml penicillin and 100 μg/mL streptomycin (cat # DE17-602E, Lonza, Basel, Switzerland), and 3% fetal bovine serum (FBS; cat # FBS-12A, Capricorn Scientific, Ebsdorfergrund, Germany).

For the cytotoxicity and plaque assays, the antiviral activity of stenoparib on SARS-CoV-2 was assessed *in vitro* using Vero E6 *Cercopithecus aethiops* kidney cells (cat # CRL-1586, ATCC, Manassas, VA, USA) maintained in Eagle’s Minimum Essential Medium (EMEM; cat # 30-2003, ATCC, Manassas, VA, USA) supplemented with 2% or 10% FBS, 100 U/mL penicillin and 100 μg/mL streptomycin (cat # P0781, Sigma Aldrich, St. Louis, MO, USA), 0.01 M HEPES solution (cat # H0887, Sigma Aldrich, St. Louis, MO, USA), 1 mM sodium pyruvate (cat # 11360070, ThermoFisher Scientific, Waltham, MA, USA), 1x non-essential amino acids solution (cat # SH3023801, Fisher Scientific, Waltham, MA, USA).

### ViroSpot reduction assay

The activities of stenoparib and remdesivir were assessed against a *wt* SARS-CoV-2 strain (BavPat1/2020), and four additional SARS-CoV-2 variants of concern; *alpha* (B.1.1.7), *beta* (B.1.351) *delta* (B.1.617.2), and *gamma* (P1), hereafter referred to using Greek nomenclature (Table 1). We performed ten, serial 2-fold dilutions of compound, mixed with 100 plaque-forming units (pfu) of virus, and added the mixture to 80% confluent Vero E6 cells growing in multi-well plates. The Vero E6 cells were fixed and stained 20 hours (h) after infection and immunostained with a SARS-CoV/SARS-CoV-2 Nucleocapsid monoclonal antibody (Sino Biological, Wayne, PA, USA) followed by a horseradish peroxidase (HRP)-labeled Goat anti-Mouse IgG (cat # G-21040, Thermo Fisher, Waltham, MA, USA) and TrueBlue Peroxidase Substrate (cat # 5510-0030, Seracare, Milford, MA, USA). Spots were counted using a CTL ImmunoSpot Image Analyzer (Cleveland, OH, USA) as previously described [10]. The EC_50_ values were approximated using package ‘drc 3.0-1’ in R version 4.1.1 [11, 12]. Percent inhibition was normalized by taking the average fraction of the wells that showed positive staining compared to the control wells with the highest positive staining [10].

**Table 1.** SARS-CoV-2 variants used in ViroSpot reduction and plaque assays to evaluate antiviral properties of stenoparib and remdesivir. The *wt*, *alpha, beta, delta,* and *gamma* strains were used in the ViroSpot reduction assay. The *alpha* strain was used in the plaque assays. The *delta* strain carries an ORF7a deletion which may have the potential to impact virulence [14]. SARS-CoV-2 strain deposition and contribution credits are provided in the Acknowledgements as requested by BEI Resources.

| Variant | Pango lineage | Strain | Isolate | Source |
| --- | --- | --- | --- | --- |
| *wt* | - | BavPat1/2020 | Germany | European Virus Archive – GLOBAL; 026V-03883 |
| *alpha* | B.1.1.7 | hCoV-19/USA/CA_CDC_5574/2020 | California, USA | BEI Resources; NR-54011 |
| *beta* | B.1.351 | hCoV-19/USA/MD-HP01542/2021 | Maryland, USA | BEI Resources; NR-55282 |
| *delta* | B.1.617.2 | hCoV-19/USA/PHC658/2021 | Tennessee, USA | BEI Resources; NR-55611* |
| *gamma* | P.1 | Isolate hCoV-19/Japan/TY7-503/2021 | Japan | BEI Resources; NR-54982 |

### Plaque assay

Vero E6 cells were infected with the SARS-CoV-2 *alpha* variant using a multiplicity of infection (MOI) of 0.1, and the activity of stenoparib was assessed with and without remdesivir using a plaque reduction assay. Following viral infection and treatment with stenoparib and/or remdesivir, cells were overlaid with low melting point agarose (cat # 1613112, BioRad, Hercules, CA, USA), fixed at 120 h post infection with 4.0% paraformaldehyde (cat # AAJ19943K2, Fisher Scientific, Waltham, MA, USA), stained with crystal violet (cat # V5265, Sigma Aldrich, St. Louis, MO, USA), and plaques were manually counted.

### Cytotoxicity Assay

Cytotoxicity was measured using the Promega CytoTox 96 Nonradioactive Cytotoxicity Assay kit (cat # G1780, Promega, Madison, WI, USA) in 50 μL reactions and 96-well format (cat # 161093, Thermo Scientific, Waltham, MA, USA) according to the manufacturer’s protocol. The assay was measured using a BioTeK Synergy HT plate reader (model # 7091000, BioTek, Winooski, VT, USA) set at 490 nm wavelength. Vero E6 cells were assessed for cytotoxicity at 5 days post-infection. Percent cytotoxicity was calculated by dividing the experimental LDH release at 490 nm wavelength (OD_490_) by the maximum LDH release control and multiplying by 100.

## Results

We explored the activity of stenoparib against a *wt* SARS-CoV-2 strain and four variant strains *alpha, beta, delta,* and *gamma* (Table 1). Inhibition of virus replication by stenoparib was dose-dependent, with 50% effective concentration (EC_50_) estimates ranging from 4.1 μM to 24.1 μM across the five virus strains tested (Fig. 1A). Interestingly, the EC_50_ estimate for stenoparib and the *beta* variant (24.1 μM) was 2.8- to 5.8-fold higher than for the *wt*, *alpha,* and *delta* variants, which were 4.1 μM, (p = 1.19×10^-4^); 8.5 μM, (p = 7.96×10^-9^); and 8.2 μM, (p = 0.06; S1 and S2 Tables). Similarly, the EC_50_ estimate for stenoparib against the *gamma* variant (13.6 μM) was lower than for the *beta* variant (24.1 μM), however this difference was not statistically significant (p = 0.22; S1 and S2 Tables). An analogous phenomenon was noted for remdesivir. The *beta* EC_50_ estimate (9.9 μM) was 2.1- to 4.3-fold higher than for the *wt*, *alpha, delta,* and *gamma* variants; 3.8 μM, (p = 2.76×10^-5^); 4.7 μM, (p = 2.05×10^-11^); 2.3 μM, (p = 4.97×10^-3^); and 4.1 μM (p = 2.32×10^-4^), respectively (S1 and S2 Tables). The ViroSpot experiment was conducted using technical quadruplicates per compound concentration, and infected and untreated cells were used as controls.

**Fig 1.**
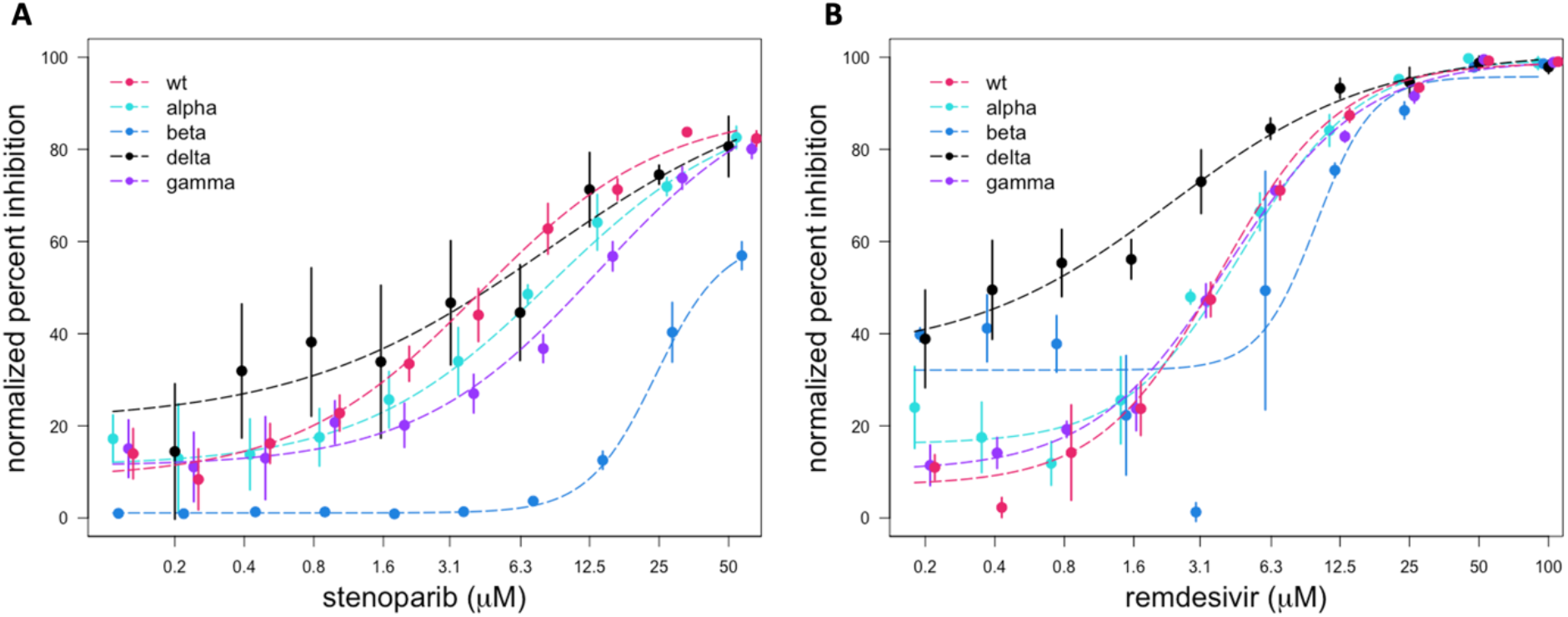
Dose-response curves for stenoparib and remdesivir on wild type SARS-CoV-2 and four variant SARS-CoV-2 strains. Normalized percent inhibition (Y-axis) of *wt* SARS-CoV-2 and variant strains *alpha*, *beta*, *delta*, and *gamma* in Vero E6 cells treated with (A) stenoparib and (B) remdesivir at the indicated concentrations (X-axis). Error bars are the standard deviation.

In an additional set of experiments, we focused on the SARS-CoV-2 *alpha* variant. Vero E6 cells were infected with the *alpha* variant using a multiplicity of infection of 0.1, and the activity of stenoparib was assessed with and without remdesivir using the plaque reduction assay. Plaques are areas of dead or destroyed cells and appear as small, clear regions in an infected cell monolayer after fixation and staining with crystal violet. We combined 2.5, 5.0, and 10 μM doses of stenoparib with the previously reported EC_50_ of remdesivir (0.5 μM) [6]. The three control types were 1) infected and untreated cells, 2) uninfected and untreated cells, and 3) infected cells treated with a combination of camostat mesylate and aloxistatin (E64d) (combination hereafter referred to as Camostat-E64d, or ‘CE’), which are protease inhibitors that prevent Spike (S) protein cleavage and virus entry into the cell [12].

As shown in Fig. 2A and 2B, neither 10 μM stenoparib nor 0.5 μM remdesivir achieved greater than a 50% reduction in plaquing efficiency compared to the infected, untreated cells. When combined, however, plaque formation was reduced by over 80% compared to infected, untreated cells. This reduction was superior to what was achievable with stenoparib (p = 3.03×10^-5^) or remdesivir (p = 1.99×10^-4^) alone at these doses. Notably, cytotoxicity remained near baseline levels (Fig. 2C). The cytotoxicity experiment was run in quadruplicate and cells were treated with CE, dimethylsulfoxide (DMSO), lysis buffer, or left untreated (no compound) as controls. When combined, the activity of two or more drugs with different mechanisms of activity may synergize, with the potential benefit of reducing individual doses of each drug and minimizing undesirable side effects in the patient.

**Fig 2.**
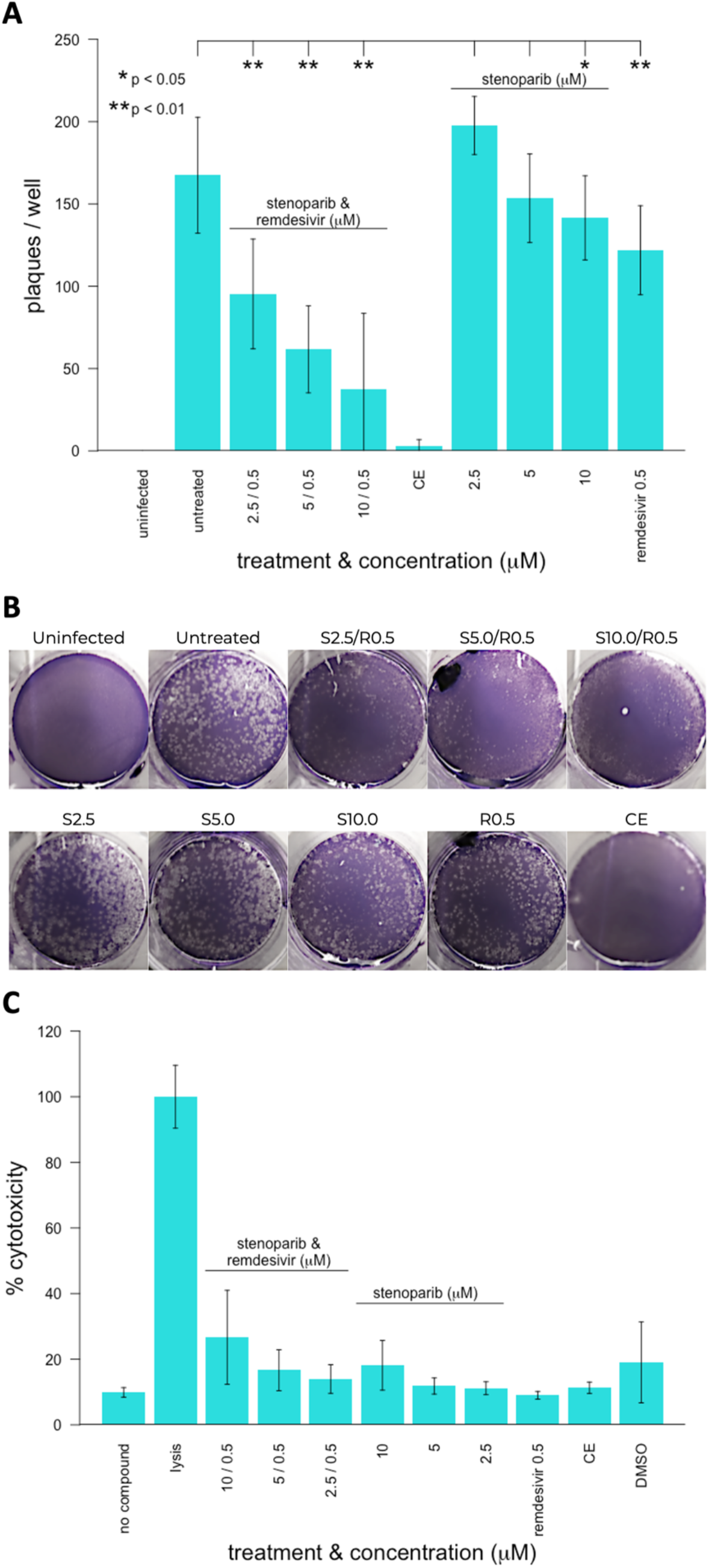
Activity of stenoparib and remdesivir on the *alpha* SARS-CoV-2 variant and cytotoxicity of stenoparib and remdesivir in Vero E6 cells. (A) Virus plaque assays. Vero E6 cells were infected for 2 hours with SARS-CoV-2 *alpha* variant, washed, and overlaid with low-melting temp agarose/MEM with or without stenoparib and/or remdesivir at the concentrations indicated. The cell monolayers were fixed with 4% paraformaldehyde and stained with 0.1% crystal violet prior to manual counting of plaques. The experiment was performed using technical triplicates and error bars represent the standard deviation of three independent experiments. For pairwise comparisons to untreated cells, * indicates p-values less than 0.05 and ** indicates p-values less than 0.01. (B) Photographs of plate wells after crystal violet staining. Labels indicate the concentrations of stenoparib or remdesivir. Abbreviations: S = stenoparib, R = remdesivir at the concentrations indicated (i.e. 2.5 / 0.5 = 2.5 μM stenoparib, 0.5 μM remdesivir). (C) Cytotoxicity measurements performed by the lactate dehydrogenase release assay (Promega Cytotox 96) 48 hours after infection. Inhibitors stenoparib or remdesivir were used at the indicated concentrations (X-axis). The assays were performed according to the manufacturer’s recommendations. Results from drug treatments were normalized to the fraction of maximum cytotoxicity (% Cytotoxicity; Y-axis) achieved by detergent lysis of cells. The samples were processed in triplicate and error bars represent the standard deviation of three independent experiments.

## Discussion

According to a recent report, the *beta* variant appears to exhibit a significantly reduced eclipse period (length of time between initial infection and the production of virus by a cell) and more rapid replication *in vitro* compared to the *alpha* variant [14], which could explain the relatively higher stenoparib and remdesivir EC_50_ estimates for the *beta* variant. In addition to numerous changes to the S protein, the *beta* variant carries some unique mutations in the N protein and in the Nsp3 polyprotein (ORF1a) that are not found in the other variants [15], and may be involved in viral degradation pathways including ADPR following infection [16]. Differential susceptibility to ADPR may be related to the altered replication kinetics of the *beta* variant, although this awaits further investigation.

Besides the known roles for the PARP1 and PARP2 enzymes in DNA repair [17, 18], members of the PARP family have numerous additional functions [19]. The 18 known human PARPs appear to differentially affect viral replication; some exhibit proviral activity and others exhibit antiviral roles [7, 20–24]. In coronavirus, the N protein is mono (ADP) ribosylated (MARylated) during infection. Stability of the N protein is critical for viral genome replication, translation, packaging, and modulation of the host cell cycle [25]. The N protein is the only ADPR target definitively identified in coronavirus thus far [20], and the prevalence of this modification across multiple coronavirus families suggests an important role in virus stability [20]. Despite this, the role of N protein MARylation, and of ADPR in general with regards to coronavirus stability, is not well characterized. Possibly ADPR has a role in regulating virus genome structure, like ADPR of histones or the adenovirus core proteins [26, 27]. While remdesivir and molnupiravir inhibit viral RNA replication, stenoparib likely acts through multiple targets. A combination of stenoparib and remdesivir or molnupiravir may be highly potent at inhibiting SARS-family coronaviruses including SARS-CoV-2.

A host-targeting therapeutic like stenoparib could be a significant benefit for COVID-19 patients as a standalone therapy, or especially as part of a combinatorial COVID-19 treatment strategy with an antiviral drug such as remdesivir or molnupiravir. Combinations of two or more drugs may lead to synergism through different mechanisms of action, which has the potential benefit of reducing individual doses of each drug and minimizing undesirable side effects.

## Author contributions

**Conceptualization**: Christopher T French, Erik W Settles, Steen Knudsen, Paul S Keim

**Data curation**: Katherine E Zarn

**Formal analysis**: Katherine E Zarn, Christopher T French, Sierra A Jaramillo

**Funding acquisition**: Steen Knudsen, Christopher T French, Paul S Keim

**Investigation:** Katherine E Zarn, Sierra A Jaramillo, Anthony R Zapata, Ashley N Jones, Haley E Nunnally, Aloys SL Tijsma, Patricia M Nuijten

**Methodology**: Christopher T French, Nathan E Stone

**Project Administration**: Christopher T French, Paul S Keim, Steen Knudsen, Aloys SL Tijsma

**Resources**: Christopher T French, Paul S Keim

**Supervision**: Christopher T French, Erik W Settles, Paul S Keim

**Visualization**: Ken Ng, Katherine E Zarn

**Writing – Original Draft**: Christopher T French

**Writing – Review & Editing**: Katherine E Zarn, Christopher T French, Nathan E Stone, Aloys SL Tijsma

## Acknowledgements

The following reagents were obtained through BEI Resources, NIAID, NIH: SARS-Related Coronavirus 2, Isolate USA/CA_CDC_5574/2020, NR-54011, deposited by the Centers for Disease Control and Prevention; SARS-Related Coronavirus 2, Isolate hCoV-19/USA/MD-HP01542/2021 (Lineage B.1.351), in Homo sapiens Lung Adenocarcinoma (Calu-3) Cells, NR-55282, contributed by Andrew S. Pekosz; SARS-Related Coronavirus 2, Isolate hCoV-19/USA/PHC658/2021 (Lineage B.1.617.2; *Delta* Variant), NR-55611, contributed by Dr. Richard Webby and Dr. Anami Patel; and SARS-Related Coronavirus 2, Isolate hCoV-19/Japan/TY7-503/2021 (Brazil P.1), NR-54982, contributed by National Institute of Infectious Diseases.

## Competing interests

SK is employed by and holds a financial interest in Allarity Therapeutics, which stands to potentially benefit from these results.

## Funding

This research was supported by a grant from The Flinn Foundation, the Cowden Endowment for Microbiology, and an Arizona Board of Regents TRIF award. Core experiments were supported by service fees paid by Allarity Therapeutics.

## Notes

### Summary of Updates

Corrected middle initials of Aloys Tijsma from SI to SL.

