## Supplemental files for "Stenoparib, an inhibitor of cellular poly (ADP-ribose) polymerases (PARPs), blocks *in vitro* replication of SARS-CoV-2 variants"

### Supporting information

**Table S1. Stenoparib and remdesivir $EC_{50}$ estimates**. Stenoparib (gold shading) and remdesivir (blue shading) $EC_{50}$ estimates for Vero E6 cells infected with SARS-CoV-2 *wt*, *alpha*, *beta*, *delta*, or *gamma* variants.

| | **remdesivir** | **stenoparib** |
|---|---|---|
| **wt** | 3.76 | 4.13 |
| ***alpha*** | 4.73 | 8.51 |
| ***beta*** | 9.87 | 24.15 |
| ***delta*** | 2.31 | 8.22 |
| ***gamma*** | 4.07 | 13.6 |

**Table S2. P-values for SARS-CoV-2 variant pairwise comparisons of stenoparib and remdesivir $EC_{50}$ estimates**. Stenoparib (above the diagonal, gold shading) and remdesivir (below the diagonal, blue shading) $EC_{50}$ pairwise comparisons for SARS-CoV-2 *wt*, *alpha*, *beta*, *delta*, or *gamma* variants.

| | **wt** | ***alpha*** | ***beta*** | ***delta*** | ***gamma*** |
|---|---|---|---|---|---|
| **wt** | | 0.12 | $1.19\times10^{-4}$ | 0.18 | 0.07 |
| ***alpha*** | 0.19 | | $7.96\times10^{-9}$ | 0.94 | 0.17 |
| ***beta*** | $2.76\times10^{-5}$ | $2.05\times10^{-11}$ | | 0.06 | 0.22 |
| ***delta*** | 0.02 | 0.07 | $4.97\times10^{-3}$ | | 0.16 |
| ***gamma*** | 0.63 | 0.41 | $2.32\times10^{-4}$ | $6.61\times10^{-3}$ | |

**stenoparib** (above the diagonal)

**remdesivir** (below the diagonal)
